# Neoepitope-specific vaccination of a patient with diffuse midline glioma targeting H3K27M induces polyclonal B and T cell responses across diverse HLA alleles

**DOI:** 10.1101/2023.04.28.538672

**Authors:** Tamara Boschert, Kristina Kromer, Taga Lerner, Katharina Lindner, Gordon Haltenhof, Chin Leng Tan, Kristine Jähne, Isabel Poschke, Lukas Bunse, Niklas Grassl, Iris Mildenberger, Katharina Sahm, Michael Platten, John M Lindner, Edward W Green

**Author notes:** Equal contribution (Tamara Boschert, Kristina Kromer, Taga Lerner). Equal contribution (John M Lindner, Edward W Green). **Corresponding authors:** Edward W. Green, John M. Lindner. **Disclosures:** TB, EWG, JML, MP, and KS are listed as inventors on a patent application describing K27M specific antibodies. MP, EWG, LB and KS are inventors of associated intellectual property related to H3K27M vaccines. MP and EWG are founders of Tcelltech GmbH. JML and TL are employees of BioMed X GmbH. JML and KK are listed as inventors on a patent application describing “System and Methods for Identifying T-cell Receptor Ligands”. **Author contributions** LB, NG, KS, and MP were responsible for clinical handling of patients and managing the associated clinical trial protocols. IP and IM managed clinical sample acquisition. IP managed PBMC immune monitoring KL and LB processed tumor, CSF, and PBMC samples, performed single cell sequencing, and performed peptide expansion assays. TB, TL, KL, KK, EWG and GH cloned and tested TCR sequences. TB, EWG and GH cloned and tested BCR sequences. CLT and TB performed single cell analyses TB, TL, JML, and EWG wrote the manuscript. MP, EWG, and JML conceptualised and directed experiments.

## Abstract

H3K27M, a driver mutation with T- and B-cell neoepitope characteristics, defines an aggressive subtype of diffuse glioma with poor survival. We functionally dissect the immune response of one patient who was treated with an H3K27M peptide vaccine and subsequently entered complete remission. The vaccine robustly expanded class II HLA-restricted peripheral H3K27M-specific T cells. Using functional assays, we characterized 34 clonally unique H3K27M-reactive T cell receptors and identified critical, conserved motifs in their CDR3 regions. Using detailed HLA mapping, we further demonstrate that diverse HLA-DQ, and -DR alleles present immunogenic H3K27M epitopes. Furthermore, we identified and profiled H3K27M-reactive B cell receptors from activated B cells in the cerebrospinal fluid. Our results uncover the breadth of the adaptive immune response against a shared clonal neoantigen across multiple HLA allelotypes and support the use of class II-restricted peptide vaccines to stimulate tumor-specific T and B cells harboring receptors with therapeutic potential.

## Main

A recurrent monoallelic, nonsynonymous mutation at amino acid position 27 in the histone-3.3 or 3.1 genes (H3K27M) defines a distinct subtype of highly aggressive diffuse midline glioma (DMG) characterized by high mortality and morbidity rates^1–3^. Due to infiltrative growth in midline brain structures and resistance to chemotherapy^4^, radiotherapy remains the main effective treatment option for DMG^5–7^. Due to a low mutational burden and a lack of immune cell infiltration, DMG is generally nonresponsive to immune checkpoint inhibition treatment but may benefit from personalized treatment or targeted immunotherapies ^5–7^. Indeed, a recent clinical trial of chimeric antigen receptor (CAR) T cell therapy targeting the uniformly upregulated disialoganglioside GD2 in DMG^8,9^ showed a radiographic improvement and reduction of clinical symptoms in three out of four patients^8^, and additional trials targeting other antigens are ongoing^9,10^.

The recurrent clonal neoepitope H3K27M represents an attractive intracellular target for immunotherapy. We and others have demonstrated presentation of H3K27M-derived epitopes on class I and class II HLA molecules capable of stimulating mutation-specific CD4^+^ and CD8^+^ T cell responses both in MHC humanized mice and in healthy human donors. Together, these studies have shown that HLA-DRB1*01:01-restricted CD4^+^ T cells and HLA-A*02:01-restricted CD8^+^ T cells induced by therapeutic long (H3K27M_p14-40_) or short (H3K27M_p26-35_) peptide vaccines, respectively, display antitumor efficacy *in vitro* and *in vivo*^11–13^.

While efficient endogenous presentation of the short epitope on HLA-A*02:01 is controversial^12–14^, a recently published phase 1 trial demonstrated that patients with H3K27M-specific CD8^+^ immunological responses to a short peptide vaccine had prolonged overall survival relative to non-responders^13^.

We and others have used long peptide vaccines targeting neoepitopes to stimulate antitumor T helper cell responses in early clinical trials^15–17^; however, the breadth of the T cell response is not well understood. Class II HLA-restricted epitopes can induce CD4^+^ T cell responses in patients with diverse HLA allelotypes^15–17^, but the precise restriction has not been assessed, mainly due to the lack of appropriate tools for high-throughput testing. Similarly, while B cell contributions have been described following long peptide vaccine administration, the quality of the antibody responses has not been assessed^15^.

We recently initiated a phase 1 first-in-human clinical trial testing the safety and efficacy of H3-vac, a long vaccine encompassing the (H3K27M_p14-40_) epitope in patients with newly diagnosed DMG (clinicaltrials.gov identifier NCT04808245). Here we present an in-depth functional study identifying and characterizing H3K27M-specific CD4^+^ T cell receptors (TCR) and B cell receptors (BCR) from the blood and CSF of a patient not eligible for the trial vaccinated with H3-vac on a compassionate use basis (Grassl et al., in revision). The patient exhibited a sustained clinical response to the vaccine, offering the unique opportunity to characterize biologically relevant immune responses. These efforts shed new light on potential mechanisms driving the survival of exceptional patients, and potentially offer new therapeutic modalities for diagnosed H3K27M^+^ DMG patients.

### H3-vac induces a CD4^+^ T cell specific immune response

Patients were vaccinated with the long H3K27M_14-40_ peptide according to a pre-defined vaccination schedule with bi-weekly administrations for 4 weeks, followed by monthly and quarterly immunizations (Grassl et al., in revision). Longitudinal blood samples were subjected to detailed immunophenotyping, including ELISpot assays and T cell receptor repertoire sequencing.

After partial tumor resection and ongoing peptide vaccination, patient ID01 showed an initial radiographic pseudoprogression followed by complete remission extending more than three years post-diagnosis. The patient displayed a strong H3K27M vaccination-induced immune response 4 weeks post vaccination (Extended Data Fig. 1). *Ex vivo* expansion of PBMCs from this timepoint in the presence of H3wt or H3K27M peptide (Fig. 1a) resulted in a general increase in IFNγ release; however, a specific increase of IFNγ-secreting cells was only observed in samples expanded and restimulated with the mutant H3K27M peptide (Fig. 1b). ELISpot assays with isolated CD4^+^ and CD8^+^ T cells from pre-expanded PBMC showed that this effect is predominantly driven by CD4^+^ T cells (Fig 1c).

**Figure 1:**
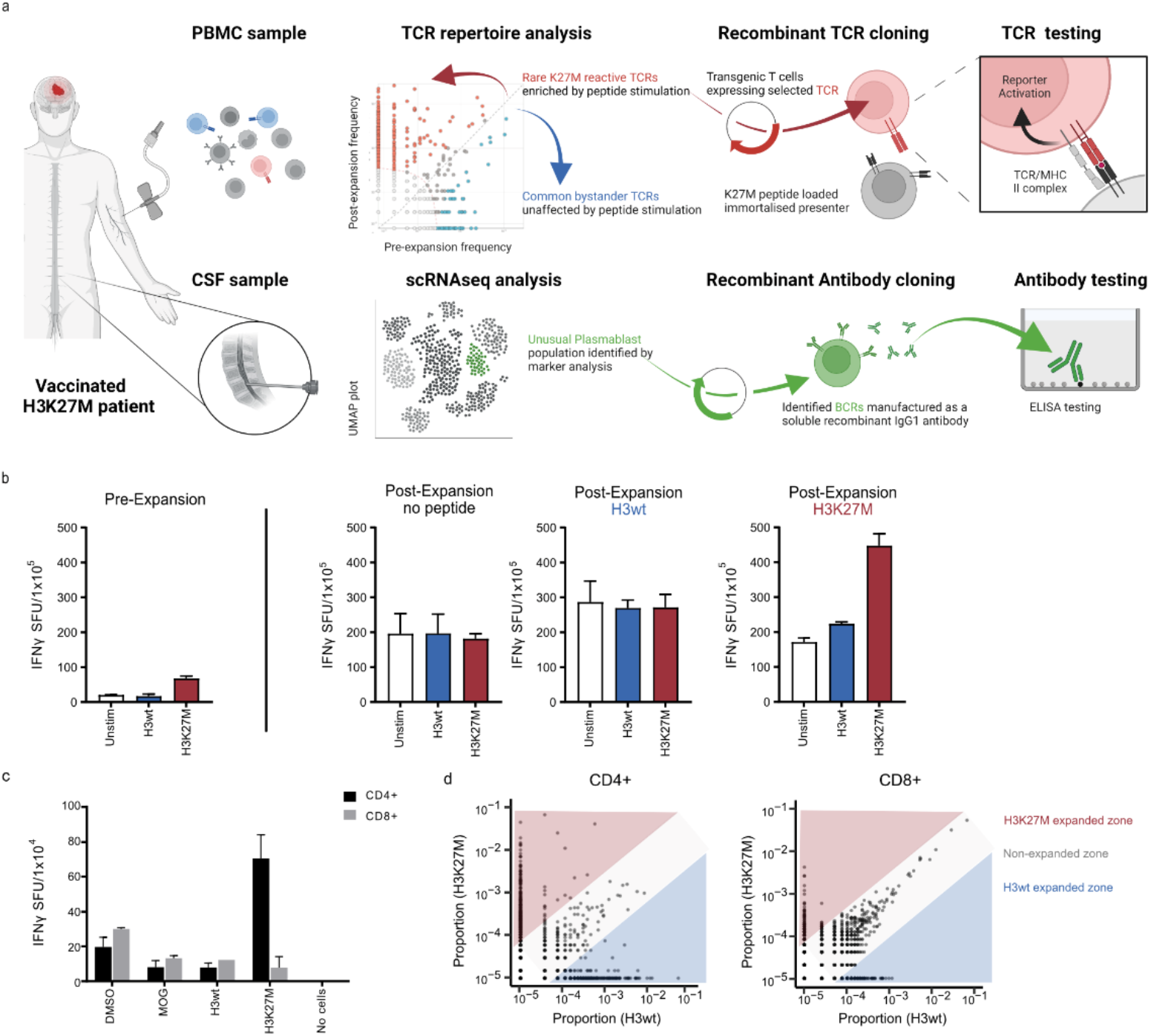
Identification and expansion of putative H3K27M reactive TCRs. **a**, Schematic of the TCR expansion, selection, and screening pipeline. Blood and CSF were collected regularly. T cells were expanded from PBMC using H3K27M peptide, H3wt peptide, or no peptide. Expansion analysis of clonotypes was performed using TCRβ deep sequencing and clonotype frequency was plotted of H3K27M expanded clones over H3wt expanded clones. Single-cell sequencing was conducted for the PBMC expanded with H3K27M peptide and CSF samples. Most frequent TCRs and BCR identified by these two selection processes are cloned and functionally validated. **b**, H3K27M specific immune response assessed by IFNγ ELISpot assay. IFNγ spots are depicted for PBMC pre-expansion and post *ex vivo* restimulation with the respective peptide. Columns represent the means of technical triplicates. Technical duplicates in panel 4. **c**, IFNγ ELISpot assay of CD4^+^ and CD8^+^ positive T cells isolated from PBMC pre-expansion. **d**, Dot plots showing the frequency of H3K27M- over H3wt-expanded clonotypes in the CD4^+^ (left panel) and CD8^+^ T (right panel) cell population post expansion.

Following peptide expansion, PBMC samples were sorted into CD4^+^ and CD8^+^ fractions and each was subjected to TCR-beta chain repertoire sequencing to assess the expansion dynamics of individual TCR clonotypes. While we observed that individual CD4^+^ T cell clonotype frequencies shifted dramatically in response to mutant or wild type H3 peptide stimulation, while CD8^+^ T cell TCR clonotypes expanded non-specifically (Fig. 1d), further suggesting that the vaccine-induced immune response was driven by CD4^+^ T cells.

### H3-vac expands a polyclonal H3K27M-specific TCR repertoire

To confirm that individual TCRs recognize H3K27M epitopes, peptide-expanded PBMCs were submitted for single-cell VDJ (scVDJ) sequencing to determine alpha/beta chain pairing and reconstitute full TCR clonotypes. Given that H3K27M-reactive TCRs effecting and/or orchestrating brain tumor clearance would be enriched in the cerebrospinal fluid (CSF), we also performed scVDJ sequencing on cells from post-treatment CSF samples. We synthesised and cloned a total of 102 TCRs, giving preference to clonotypes with the greatest *ex vivo* expansion, with the expectation that these were most likely to be H3K27M-specific. TCR clonotypes expressing two alpha chains were screened separately but considered to be representative of a single *bona fide* TCR clone (Fig. 2a, Extended Data Fig. 2d, Extended Data Table 1).

**Figure 2:**
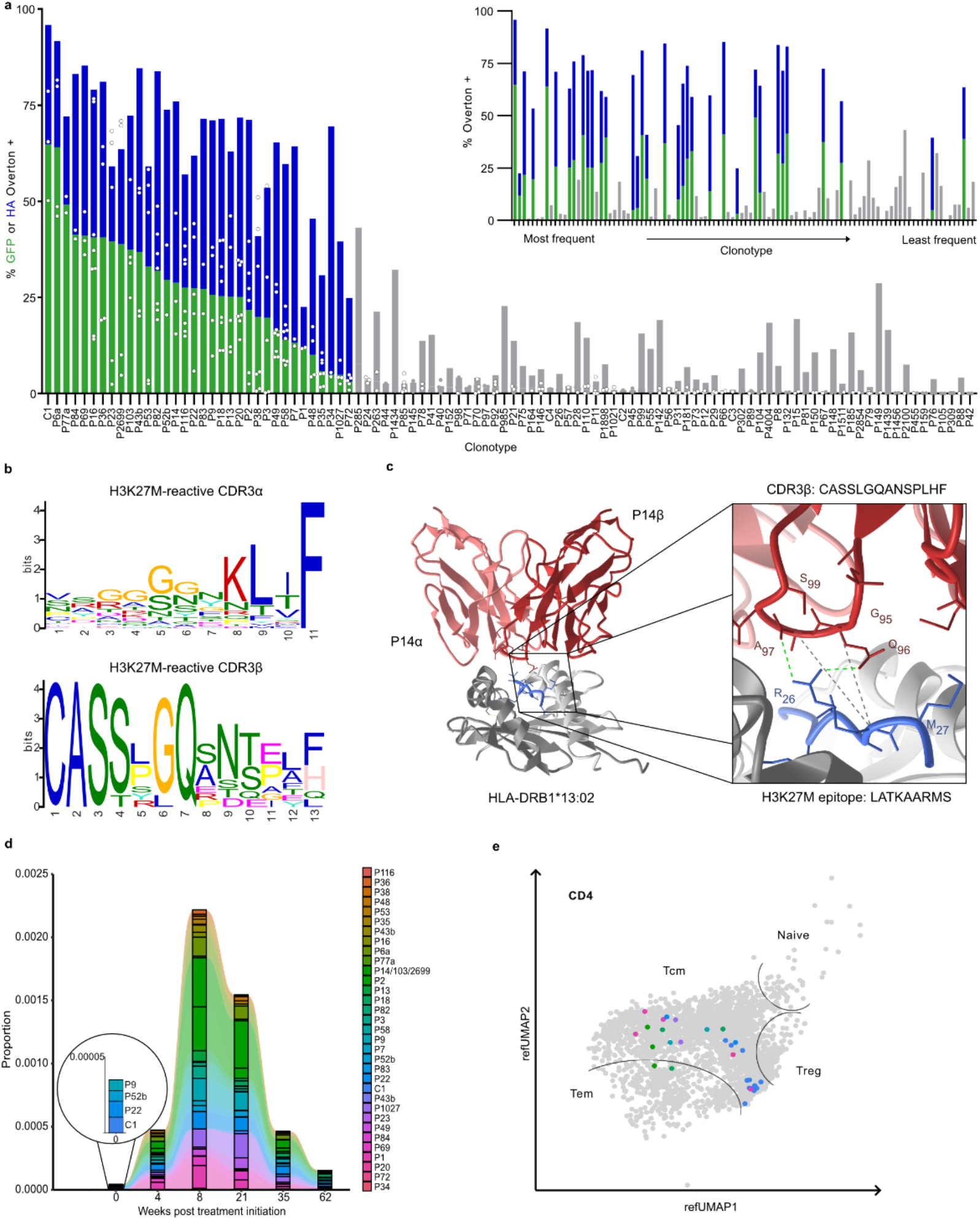
Validating the H3K27M reactivity of patient ID01 TCRs in a Jurkat reporter line. TCR nomenclature in figures reflects their origin and frequency, P indicates the TCR was identified first from expanded PBMC and C from the CSF, with the index indicating their frequency rank. The lower the number, the more frequent the TCR. If a beta chain paired with two alpha chains shares the same frequency, this is reflected by TCRs named with the same clonotype number followed by a or b, e.g. P6a/ P6b. **a**, Superimposed dual reporter signals from GFP (green) and HA staining (blue) after co-culture of transduced reporter cells and ID01 B cell line with 10µM H3K27M peptide. The signal is determined by flow cytometry using Overton positive referenced to H3wt treated samples. TCRs are arranged by GFP signal (main plot) or clonotype frequency (upper right corner). Gray TCRs do not pass criteria to be considered reactive (Extended Data Fig. 3). **b**, Amino acid motif of CDR3 alpha (upper panel) and beta (lower panel) chains of all reactive TCRs. Motif analysis was performed using XSTREME tool. Shuffled input sequences and TCRb deep sequencing data of the baseline repertoire were used as control sequences for finding the CDR3α and CDR3β motif, respectively. **c**, High-resolution modelling of complete pMHC:TCR complex (left panel) of H3K27M reactive TCR P14. Interacting bonds of the CDR3β (red) residues with the peptide (blue) are indicated in the magnified panel. **d**, Longitudinal TCRb deep sequencing was performed to track the proportion of H3K27M reactive clonotypes in peripheral blood at the indicated timepoints. **e**, UMAP of CD4^+^ T cell cluster indicating T cell identity identify of reactive TCR clonotypes (right panel) from CSF aspirates 35 weeks after treatment initiation.

TCRs were screened for reactivity against H3K27M using the T-FINDER platform (Schmid and Cetin et al, in preparation). Briefly, patient-derived B cells were immortalized to generate a fully autologous and HLA-competent B lymphoblastoid cell line (B-LCL), while Jurkat cells with an integrated sensitive dual reporter of T cell activation were transgenically modified to express each TCR of interest (Extended Data 2a). The B-LCLs were pulsed with H3wt or H3K27M peptides and co-cultured for 16h with each transgenic TCR reporter cell line. Upon cognate TCR:peptide interactions, the reporter produces a T cell-intrinsic GFP signal and marks co-cultured B-LCLs by secreting an HA-tagged αCD19 scFv, both of which can be assessed by flow cytometry.

Of the 102 TCRs from patient ID01 that were tested, 34 reacted specifically to the H3K27M mutant peptide but not the wild-type control (Fig. 2a). Although several less frequent TCRs also showed H3K27M reactivity, those with the strongest *ex vivo* expansion were indeed enriched for reactive clonotypes (Fig. 2a inset panel). For clonotypes natively expressing two alpha chains, in all cases only a single alpha chain was reactive, demonstrating that binding of the H3K27M epitope required contributions from both chains (Extended Data Fig. 2e).

TCRs varied dramatically in their response to H3K27M; we assigned a series of conservative thresholds to eliminate false positives, *e*.*g*. excluding TCRs with cross-reactivity to H3wt (see Extended Data Fig. 2b for further details). Specific H3K27M TCR activities ranged from 64.8 % GFP positive / 95.9 % HA positive to 3.1 % GFP positive / 24.9 % HA positive. The TCR with the greatest activity in our co-culture assay was also the most prevalent TCR in the CSF (C1); the low frequency of this sequence prior to vaccination is indicative of strong clonal expansion and selection *in vivo* (Fig. 2d). Overall, patient ID01’s tumor-specific TCR repertoire was highly polyclonal (Extended Data Fig. 2c), though we did observe evidence of convergent CDR3 selection: TCRs P14, P103, and P2699 all share the same CDR3α and CDR3β amino acid sequence but differ at their recombination junction site nucleotide sequences (Extended Data Fig. 2d). This phenomenon has previously been associated with antigen-specific T cell responses to cancer immunotherapies^18^.

Relative to baseline, a longitudinal analysis of TRBV and TRBJ gene usage across the entire PBMC TCR repertoire shows no changes between visits (Extended Data Fig. 3a). However, the presence of TRBV7-2 in patient ID01 is significantly overrepresented in the reactive TCRs relative to the baseline TCR repertoire (Fisher’s exact test, p < 0.005) (Extended Data Fig. 3b). Additionally, we analyzed the recombination and usage of TRBD gene segments after observing the potential overrepresentation of a central glutamine residue in the CDR3β regions of reactive TCRs. The human TRBD locus contains two gene segments that can be inserted directly or inversely and read in three translational frames; only one combination of gene segment selection, orientation, and reading frame encodes this glutamine (Q) residue. Relative to the baseline TCR repertoire, the appearance of glutamine among reactive TCRs is, indeed, statistically enriched (Fisher’s exact test, p < 0.00005) (Fig. 2b). In fact, this glutamine residue forms the core of an enriched sequence motif present in the CDR3β of 8 peptide-reactive TCRs, which is frequently observed in combination with a significantly enriched GGx_1-3_K motif in cognate TCR CDR3α sequences (Fig. 2b). To better understand the relevance of these motifs, we applied structural TCR-pMHC modeling using ImmuneScape^19^. While structural and/or dynamic modelling studies are required to fully characterize these interactions, the CDR3β-conserved glutamine appears to interact directly with R_26_ of the H3K27M peptide in a model of the P14 TCR in complex with the peptide LATKAARMS presented by HLA-DRB1*13:02 (Fig. 2c, see the next section of results for TCR restriction mapping).

Cataloging this set of functionally validated H3K27M peptide-reactive TCRs enabled a focused re-analysis of the longitudinal TCR sequencing datasets. While reactive clonotypes could not be identified using TCR network analysis only, a *post hoc* analysis revealed significant expansion of H3K27M-reactive clones within the full repertoire for 8 weeks after administration of the first vaccine, followed by a decrease (Fig. 2d). At baseline, P52b, P9, P22, and C1 were detectable at very low proportions in bulk TCRβ sequencing. At 62 weeks post-treatment initiation, the proportion of reactive TCRs remained higher than at baseline. At weeks 8 and 21, when the clonal expansion was at its greatest, the most frequent TCRs in the peripheral blood were the clonally converged P14/P103/P2699. At the final time point, these TCRs also comprise the greatest portion of reactive TCRs (Fig. 2d). A subset of H3K27M-reactive TCRs was also found in the CSF (Fig. 2e), including the convergently selected P14/P103/P2699 cluster, as well as P1, P2, P9, P18, P20, P22, P69, and P1027. We complemented our scVDJ with scRNA sequencing and observed that these T cells almost exclusively exhibited a CD62L^+^CCR7^+^CD27^+^CD95^+^TCF7^+^ central memory phenotype (Fig. 2e).

### H3K27M epitopes are presented by both HLA-DR and HLA-DQ alleles in patient ID01

The diversity of HLA alleles which can present H3K27M-derived epitopes critically determines the number of patients that would benefit from therapeutic vaccination and/or TCR-T cell therapies. Computational algorithms such as NetMHCIIpan do not predict any strongly binding HLA:epitope complexes^20^; therefore, we applied a two-step approach to empirically determine the HLA restriction (and thus the peptide-presenting allele) of each reactive TCR.

First, we generated HLA-DPA, HLA-DQA, and HLA-DRA knockouts of patient ID01’s B-LCL (Extended Data Fig. 4a). The resulting cell lines were loaded with either H3K27M or H3wt peptide and co-cultured with reactive TCR-transgenic reporter cells (Fig. 3a-c, Extended Data Fig. 2a). As expected, disrupting the restricting HLA locus abrogated TCR reactivity completely, with most clonotypes showing restriction to HLA-DR and several to HLA-DQ (Fig. 3a,b).

**Figure 3:**
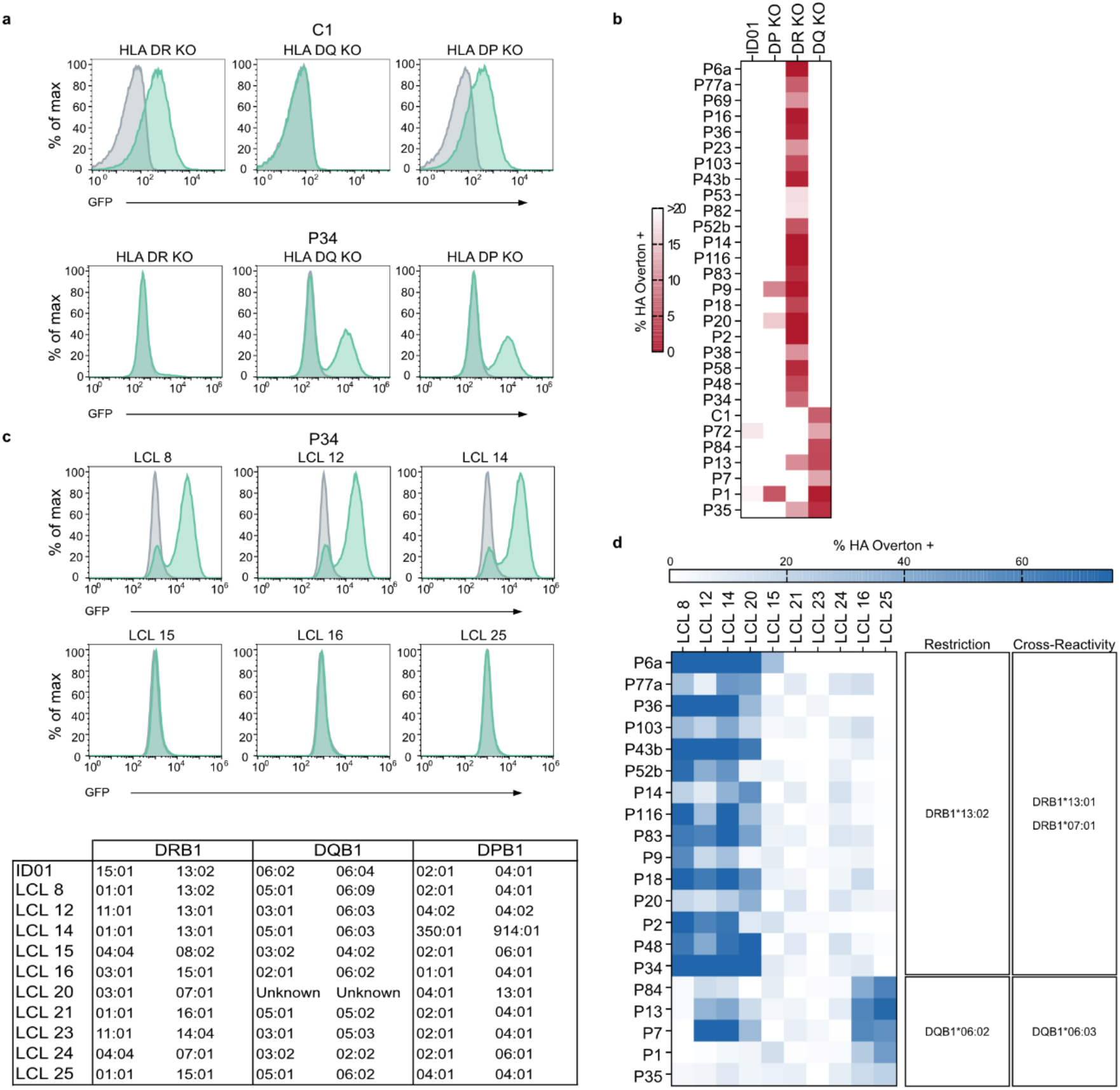
H3K27M HLA restriction analysis. **a**, Representative flow cytometry histograms of H3K27M reactive TCRs co-cultured with peptide-pulsed autologous HLA class II knock-out B-LCL (H3K27M in green or H3wt peptide in grey) depicting GFP reporter levels. Heatmaps of H3K27M peptide reactive TCRs co-cultured with either autologous donor B-LCL or their corresponding HLA class II knock-outs of **b**, ID01. **c**, Top, representative histograms of HLA allele restriction screening against B-LCL from a healthy donor library. Bottom, HLA alleles of ID01 and LCL lines used in analysis. **d**, Restriction analysis screening heatmap of H3K27M reactive TCRs from ID01. Percentage of HA positive B-LCL was determined by the Overton method using H3wt peptide-pulsed controls for histogram subtraction.

In the second step, a panel of healthy donor B-LCL that share one or more HLA alleles with ID01 (Fig. 3c, see Extended Data Table 2 for detailed HLA types) was used to identify the restricting HLA allele and screen for additional, cross-restricted HLA alleles. The healthy donor B-LCL were peptide pulsed and co-cultured with reactive TCR-expressing T cells as above, and the combination of response-inducing lines was used to identify the presenting allele of each TCR (Fig. 3d). All ID01 HLA-DR responsive TCRs were positive with LCL 8, LCL 12, and LCL 14, indicating they are restricted to DRB1*13:02, as well as the closely related non-autologous DRB1*13:01 allele. Most of the HLA-DR restricted TCRs also showed some response to peptide presentation by lines LCL 24 and LCL 20, indicating that the H3K27M peptide is presented by DRB1*07:01 – contrary to HLA supertype predictions^21^. The five HLA-DQ restricted TCRs were mapped to DQB1*06:02 (LCL lines 16 and 250), and with the exception of the weakly reactive TCR P1, also reacted with the DQB1*06:03-expressing LCL lines 12 and 14.

### Patient CSF contains a clonally expanded, H3K27M-reactive B cell population

Having identified tumor-reactive TCRs in the CSF of patient ID01, we compared the composition of this sample to publicly available control datasets^22^ using Azimuth to map data onto a common reference^23^. We observed prominent groups of CD19^+^CD20^+^CD27^lo^IgD^-^CD38^-^ CD138^-^ memory B cells and CD19^-^CD20^-^CD27^+^IgD^-^CD38^hi^CD138^hi^ plasmablasts in patient ID01 but not in healthy donors (Fig. 4 a,b, Extended Data Fig. 5). The BCR sequences of these cells showed strong pauciclonality: the 5 most frequent clones comprise approximately 46% of the overall repertoire (Fig. 4c, n=80 cells). We tested the two most frequent BCRs (9.5% and 7.6% respectively, Fig. 4c) for their H3K27M binding capacity by producing both as recombinant IgG1 antibodies. We used direct ELISA to show binding specifically to the H3K27M peptide with similar functional avidities (EC_50_ BCR1 13.1 nM, BCR2 16.5 nM) (Fig. 4d). To determine whether these antibodies bind full-length H3K27M as well as the vaccinating 27-mer H3K27M_p14-40_ peptide, we assessed their affinity in competition with the full-length H3K27M and H3wt proteins. Both antibodies displayed high affinity to the H3K27M protein and the peptide within a range of 10^−7^ and 10^−8^ M, respectively (Fig. 4e). Neither antibody bound appreciably to the H3wt peptide at physiologically relevant concentrations.

**Figure 4:**
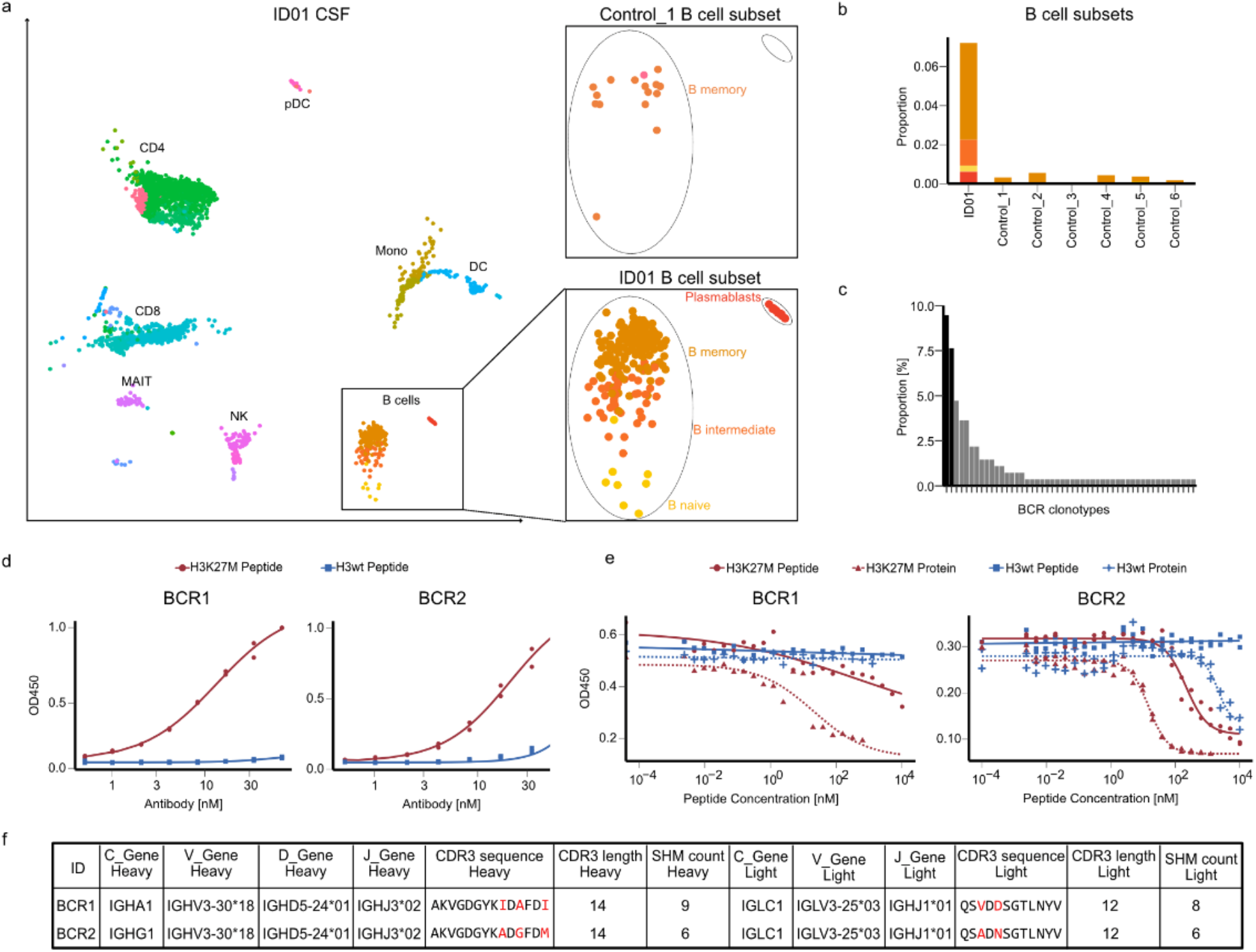
Analysis of cellular composition of ID01 CSF. **a**, Single cell sequencing data of ID01 CSF (n = 1 sample, n = 5,316 single cells, left) projected as UMAP. Immune cell clusters were identified by applying Seurat v4 reference mapping. Magnified panels show B cell subpopulations of ID01 compared to a Control CSF sample. **b**, Relative abundance of B cell subclasses in ID01 and control CSF. **c**, Histogram depicting frequency of identified BCRs in the CSF. Tested BCRs are highlighted in black. **d**, Dose response curves of BCRs were assessed using a direct ELISA pre-coated with either H3wt or H3K27M peptide. OD was measured at 450 and plotted over antibody concentration in nM. Technical duplicates in both panels. **e**, Affinity to the H3K27M peptide was determined in a competitive ELISA in which the BCRs were incubated with the respective peptide or protein overnight and then added to H3K27M precoated ELISA plates. Dose response curves show the OD450 over peptide concentration in nM. **f**, Table depicting the VDJ gene usage, CDR3 amino acid sequences and their acquired somatic hypermutations (SHM) of both chains of BCR1 and BCR2. Highlighted amino acids indicate differences in their CDR3 amino acid sequences.

Analysis of the heavy and light chain sequences of these two BCRs shows they share a common clonal origin (Fig. 4f). Both receptors exhibit significant somatic hypermutation, indicating a history of T-cell supported affinity maturation from their germline sequences; furthermore, single-cell transcript data showed that both BCR1 and BCR2 had undergone class-switch recombination to the IgA and IgG1 isotypes, respectively. These results highlight the highly functional CD4^+^ T cell response induced by the peptide vaccine, since cognate B-cell differentiation is driven by CD4^+^ T cells recognizing the same epitope^24^.

## Discussion

In this study, we investigated the immune response of a patient vaccinated with a neo-epitope-containing peptide and conducted a comprehensive functional analysis of putative H3K27M-cognate TCRs. We demonstrated that vaccination results in a robust CD4^+^ T cell response leading to a large, polyclonal repertoire of neoantigen-specific TCRs and targeted activation of H3K27M-specific B cells in the CSF.

Furthermore, we demonstrated that the H3K27M peptide vaccine induces an immune response across diverse HLA alleleotypes: the deconvoluted HLA restrictions of cognate CD4^+^ TCRs demonstrate restrictions to members of multiple class II HLA loci. DRB1*13:02 and DQB1*06:03 frequently co-occur on the same haplotype, whereas DRB1*07:01 and DQB1*06 are not a common haplotype pair^25^. Considering these allelic frequencies and the TCRs described in this work with their restrictions, 40-45% of the German population (where the INTERCEPT-H3 trial – NCT04808245 - is currently ongoing) could be covered with cell therapy using a pair of identified TCRs. It is notable that many epitope-specific TCRs analyzed here are restricted on the DRB1*13:02 allele, despite this allele:peptide combination not being clearly predicted using common *in silico* methods. This observation suggests that prediction-based approaches must continue to be supported by functional studies, at a minimum to optimize machine learning or other algorithms used to make such predictions in order to reduce the number of false negatives in future computational work.

The presence of convergently recombined TCRs (i.e., TCRs that have a shared antigen specificity and identical amino acid sequence but unique nucleotide sequences) has been proposed as a biomarker and indicator for cancer immunotherapy responses^18,26,27^. While additional clinical validation of convergent TCRs as a marker is required, it is of great interest that such TCRs (P14, P103, and P2699) are detected in ID01. The use of UMIs during reverse transcription, the number of corroborating sequencing reads, and the position of divergent nucleotides directly at the V(D)J recombination junctions strongly suggest that these TCRs are indeed convergent clonotypes. Furthermore, these TCRs are highly abundant and occupy a significant clonal niche (Fig 2d), and their relevance is highlighted by their presence within the CSF (Fig 2e).

We have shown the presence of increasing H3K27M-reactive TCR frequencies in the CSF of patient ID01 upon subsequent vaccine administrations (Fig 2e). It is striking that T and B cells with neoantigen-cognate receptors would home to the CSF following a subcutaneously administered peptide vaccine unless they detect the endogenous epitope within the central nervous system, presented either by tumor cells or scavenger cells such as macrophages or dendritic cells that have taken up H3K27M-containing debris following tumor cell death. Unfortunately, we were not able to confirm migration into the tumor due to limited sample availability because of the midline and therefore critical localization of these tumors. However, an in-depth study on the immune cell repertoire of matched brain metastases and CSF has shown consistently high clonal overlap of TCRs, suggesting immune cell trafficking between these compartments^28^. The ability of immune cells to migrate between the CSF and brain lesions was also confirmed in studies focusing on non-oncogenic lesions^29^.

The H3K27M-reactive TCRs in the CSF were identified as central memory CD4^+^ T cells, which have previously been shown to be superior in controlling tumor growth both *in vitro* and *in vivo* due to their increased cytokine secretion and prolonged persistence relative to effector T cells^30^. Moreover, patients successfully treated with immunotherapy maintain highly functional memory T cells years after treatment^31^. The functionality of the CD4^+^ T cell clones identified in this study is further underlined by the prominent cluster of activated B cells expressing H3K27M-specific BCRs in the CSF. The presence of such tumor-reactive B cells and their intrinsic ability to process and present antigens may support the maintenance of a functional memory CD4^+^ T cell population^32^. Interestingly, the formation of B cells in tertiary lymphoid structures as well as neoantigen-targeted antibodies have been shown to be correlated with responsiveness to immunotherapy^33–35^. Although not yet explored in detail, genetically engineered B cells carrying tumor-reactive BCRs may comprise another potential immunotherapeutic strategy^36^. In addition to their beneficial effects on CD4^+^ T cells, they could also indirectly promote tumor killing by activating the complement system upon secretion of neoantigen-specific antibodies, which may recognize HLA-bound neo-epitopes *in vivo*^36^.

In concordance with previous publications exploring immune responses following neoantigen vaccination^11,16,37^, we show that CD4^+^ T cells are the chiefly responding subset in the vaccinated patient ID01. Often overlooked, there is accumulating evidence suggesting that CD4^+^ T cells are at least as critical as cytotoxic CD8^+^ T cells for efficient tumor clearance^38,39^. We have recently published that functional activation of CD4^+^ T cells by glioma-infiltrating myeloid cells is a major determinant of CD8^+^ T cell fitness and prevents their differentiation into terminally exhausted T cells^40^. Additionally, we found this to be critical for responsiveness to immune checkpoint inhibition. Due to the unavailability of a primary tumor cell line for patient ID01, we did not have the opportunity to functionally validate the TCRs in an autologous setting. However, the observed early-onset pseudoprogression as well as a positive proximity ligation assay, suggest a relevant and functional CD4^+^ T cell response *in vivo* against the primary tumor presenting the neoepitope H3K27M on HLA-DR molecules.

Few vaccination response studies consider the functional immune receptor repertoire. By performing such studies, it may be possible to learn about the nature of target-specific TCRs, potentially enabling their direct identification by streamlined repertoire analysis in the future^41^. If no such pattern can be identified, functional analysis will remain critical, since the sequenced TCR repertoire (even following *ex vivo* antigen-specific enrichment) is clearly not equivalent to the functional repertoire.

This detailed exploration of the remarkable and long-lasting immune response associated with long-term survival of patient ID01 enabled the identification of TCR and BCR sequences that can both be engineered to create additional cell-based therapies. With the expansion of peptide vaccination to the INTERCEPT-H3 clinical trial cohort, further data will help identify both patient factors leading to responses similar to those described here and individuals most likely to benefit from this precision treatment. Finally, the insights gained from characterizing immunotherapeutic responses in DMG are a potent proof-of-concept for both peptide vaccinations as a therapeutic strategy, as well as leveraging immunogenic T and B cell responses into off-the-shelf, personalized biologic and cellular therapies for further tumor types in the future.

## Methods

### H3K27M vaccination

The 27-mer H3K27M peptide (p14-40) was synthesized by the good manufacturing practice (GMP) facility of the University of Tübingen, Germany and 300 µg was emulsified in Montanide (ISA50)^34^ at the University Hospital Heidelberg and Mannheim, Germany within 24h before vaccination as described previously^42^. H3K27M was administered subcutaneously with subsequent injection of imiquimod (5% Aldara). Patients received H3K27M-vac after providing written, signed informed consent at the University Hospital of Mannheim and treatment was approved by the institutional review board.

### PBMC isolation

Heparinized blood from patients was diluted with phosphate-buffered saline (PBS) and layered onto Biocoll Separation Solution (Biochrom) in Leucosep tubes (Greiner Bio-One) followed by a density-gradient centrifugation (800g without brake at room temperature). Until further analyses PBMC were frozen in 50 % freezing medium A (60% X-Vivo 20, 40% fetal calf serum (FCS)) and 50% medium B (80% FCS, 20% DMSO) and stored in liquid nitrogen at −140 °C.

### IFNγ ELISpot of PBMC

White-bottom ELISpot HTS plates (MSIPS4W10, Millipore) were hydrophilized with 35 % EtOH and subsequently coated with anti-human IFNγ (1-D1K, Mabtech). Blocking was performed with X-Vivo-20 (Lonza) supplemented with 1 % human albumin (Sigma). PBMC were thawed and rested overnight in X-Vivo-20. Per well, 4 × 10^5^ cells were seeded and stimulated with 2 µg peptide in a total volume of 100 µl. H3K27M (p14-40), H3wt (p14-40) or MOG (p35-55) were used as stimulants and 10 % DMSO diluted in aqua ad iniectabilia (Braun) at equal volume was added as negative control.Positive controls were 1 μg staphylococcal enterotoxin B (Sigma-Aldrich) and 0.05 μg CMV with 0.05 μg AdV per well. After 44 h, detection of IFNγ-secreting cells was performed by adding biotinylated anti-human IFNγ antibodies (7-B6-1), streptavidin-ALP (both Mabtech) and ALP colour development buffer (Bio-Rad). IFNγ Spots were quantified by an ImmunoSpot Analyzer (ImmunoSpot/CTL Europe).

### Peptide specific T cell expansion

PBMCs were thawed and rested overnight in X-Vivo 20 medium. The next day, cells were resuspended in fresh X-Vivo 20 + 2% human albumin and counted. Half of the available PBMCs were plated in one 24-well plate at 0.5 × 10^6^ cells per 500 µL per well and placed in a 37°C CO_2_ incubator. The remaining cells were plated at the same density in a second 24-well plate but pulsed with (I) 2 µg long H3K27M peptide (p14-40), (II) 2 µg long H3wt peptide (p14-40), or (III) no peptide to control for unspecific expansion, and also placed in the incubator. After 4 hours, unpulsed cells were added to peptide pulsed wells to a final density of 1 × 10^6^ cells per well.

On day 4, 7, 9 and 11, half of the medium was replaced by 500 µL fresh medium containing 100 IU/mL IL-2 (Novartis), 50 ng/mL IL-7 (Miltenyi) and 50 ng/mL IL-15 (Miltenyi). On day 14, cells were harvested and rested overnight in cytokine-free media. The next day, cells were subjected to IFNγ ELISpot analysis as described above to verify the peptide-specific expansion of T cells. 5 × 10^4^ cells were plated per well and restimulated for 44 h with 10 µg/mL H3K27M peptide, H3wt peptide, left unstimulated or exposed to PMA/ionomycine (20 ng/mL and 1 µg/mL), respectively. Viable cells were cryopreserved for single cell sequencing.

Expanded cells were also subjected to FACS-based sorting of CD4^+^ and CD8^+^ T cell populations using a BD FACS Aria Fusion I Sorter. Briefly, cells were stained with fixable viability dye (AF700, eBioscience) and then stained with fluorescently labelled antibodies: anit-CD3-BV510 (clone HIT3A, BD), anti-CD4-BV786 (clone SK3=Leu3a, BD) and anti-CD8a-PerCP/Cy5.5 (clone RPA-T8, Invitrogen). After staining, CD4^+^ and CD8^+^ cell populations were sorted into 0.04 % BSA in PBS using a 100 µm nozzle. After FACS sorting, cells were spun down and pellets of CD4^+^ and CD8^+^ enriched cell populations were used for TCRβ deep sequencing.

### TCR beta repertoire deep sequencing

For TCR beta chain (TCRb) deep sequencing genomic DNA was isolated from PBMC or CD4^+^ and CD8^+^ sorted T cell populations following the DNeasy Blood and Tissue Kit (Qiagen) protocol. Subsequent library preparation was performed by using HsTRBC Kit V7 (Adaptive Biotechnologies) following the manufacturer’s protocol. Libraries were sequenced on an Illumina MiSeq by the Genomics & Proteomics Core Facility, German Cancer Research Center (DKFZ). The provided platform by Adaptive Biotechnologies was used for data processing. Data analysis and visualization was performed using the immunarch 0.6.6 package in RStudio.

### Single-cell RNA and VDJ sequencing

Library construction of the CSF sample was generated using Chromium Single Cell V(D)J Reagent kit v1.1 chemistry (10x Genomics; PN-1000006, PN-1000020, PN-1000005, PN-120262) following the manufacturer’s protocol. The constructed scVDJ library and scGEX libraries were sequenced on a NovaSeq6000 platform (Illumina) respectively. Control CSF datasets were retrieved from GEO with the accession code GSM4104122. Raw sequencing data was processed running cellranger pipeline (version 7.0.0) applying default settings. Subsequent analyses including quality control were performed with Seurat v4^43^. The multimodal PBMC reference dataset^23^ was used to identify cell clusters and was applied using the default settings of FindTransferAnchors() and MapQuery(). The package ScRepertoire v1.5.4 was used to integrate scVDJ and scGEX datasets. pMHC::TCR modelling was performed using ImmuneScape^19^.

### B cell immortalization

B cell isolation was performed using the the EasySep Human B Cell Isolation Kit (StemCell Technologies). PBMC were diluted in PBS supplemented with 2 % FBS and 1 mM EDTA to a final concentration of 10^7^ cells/ml. Isolated B cells were resuspended in B-LCL-medium containing 2.5 μg/ml CpG ODN 2006 (InvivoGen) and 30 μg/ml holo-transferrin (Sigma-Aldrich). Previously quantified Epstein Barr Virus (EBV) - containing supernatant from B-95 cells was added to the cell suspension before plating 5 × 10^5^ cells per well in a round-bottom 96-well plate.

### Cell culture

B-LCL were cultured in RMPI 1640 Medium (Gibco) supplemented with 10% FBS (PAN-Biotech), 1% penicillin-streptomycin (Capricorn Scientific), 50 mM β-mercaptoethanol, 1 mM sodium pyruvate (Thermo Fisher Scientific) and 5ml MEM Non-Essential Amino Acids (Gibco). Jurkat T cells (Leibnitz Institute DSMZ #ACC282) were cultured in RPMI 1640 Mediums supplemented with 10% heat-inactivated FBS and 1% penicillin-streptomycin. HEK293FT (Thermo Fisher #R70007) cells were cultured in DMEM (Gibco) supplemented with 10% FBS.

### TCR cloning and transgenesis

Sequences for TCR α-/β-variant domains were assembled into a fully human lentiviral TCR-expression vector with TCR α- and β 23 constant domains. TCR-expressing plasmids were obtained from TWIST Biosciences. Lentivirus was produced in HEK293T cells in DMEM medium. HEK cells were transfected with TCR expression vector and lentiviral transfer plasmids using TransIT-VirusGen Transfection Reagent (Mirus #MIR6700) following the manufacturer’s recommendation. Viral supernatant was harvested after 48 h and sterile filtered with a 0.45 µM syringe filter.

### Co-culture assays

B-LCL and TCR-transgenic Jurkat T cells were plated in a 1:3 cell ratio in a round-bottom 96-well plate with a final concentration of 10 µM peptide. Co-cultures were analysed after 16 h of incubation.

### TCR reactivity testing by flow cytometry

For assessing TCR reactivity, co-cultures were stained with antibodies targeting surface proteins: anti-CD3-PE/Cy7 (clone HIT3A), anti-CD20-PE/Cy7 (clone 2H7) or anti-CD20-BUV395(clone L27, BD Biosciences), anti-CD2-PerCP/Cy5.5 (clone RPA-2.10), anti-HA-APC (clone 16B12) and anti-CD69-APC/Cy7 (clone FN50) (all BioLegend). Data was acquired on an BD FACS Aria II and a Sony ID7000 Spectral Analyzer.

Data analysis for all experiments was performed using FlowJo software v.10.8.1.

Determination of TCR reactivity is described in detail in Supplementary Figure 2.

### Generation of HLA knock-out lines

HLA class II knock-out lines were generated using the published gRNA sequences from Lee et al ^44^. The gRNA were ordered as crRNA and combined in an equimolar ratio with Alt-R® CRISPR-Cas9 tracrRNA (IDT). Prior to electroporation (Neon Transfection system, ThermoFisher Scientific), Alt-R guideRNA duplex was incubated with the Alt-R® S.p. HiFi Cas9 Nuclease V3 (IDT) to form a ribonucleoprotein (RNP) complex. Electroporation was performed with Neon 10 µl tips. Briefly, 5 ×10^5^ cells were taken up in 9 µl R-Buffer containing 1 µl of RNP complex and 2 µl of Alt-R® Cas9 Electroporation Enhancer (IDT) and electroporated with one pulse at 1400 V for 30 ms. The cells were transferred into 1 ml of B-LCL medium and expression of HLA-DRA, HLA-DQA and HLA-DPA was analysed 6 days later. Negative populations were sorted with a BD FACS Aria II.

### BCR cloning

Sequences for BCR heavy and light chains were assembled into the respective ‘AbVec’ heavy and lambda chain expression vectors^45^, and prepared as endotoxin-free midipreps using the ZymoPURE II Plasmid Midiprep Kit (Zymo Research)

### Antibody production and purification

Antibodies were produced in HEK293FT cells in DMEM medium supplemented with 2 % FCS. HEK cells were transfected with both heavy and light chain BCR constructs using FuGENE HD transfection reagent (Promega) in a 1:3 ratio. Antibody-containing supernatant was harvested after 5 days and filtered through a 0.45 µM syringe filter. Antibody purification was performed using MabSelect (Cytiva) according to the manufacturer’s guidelines.

### IgG ELISA

Indirect ELISA was performed in MaxiSorp plates (Nunc) pre-coated with human H3K27M and H3wt (p14-40) (180 ng per well in PBS). Purified antibodies were added in serial dilutions.

For performance of the competitive ELISA, 96-well polypropylene plates (Greiner) were blocked with 100 % FCS for 2 h at room temperature. 125 ng of antibody was diluted with H3K27M and H3wt peptide and protein, respectively and incubated overnight at 4°C. Peptide and proteins were 1:2 serial diluted starting from 144 ng. The following morning the mixture was added to a H3K27M (p14-40, 180 ng) pre-coated MaxiSorp plate.

Plates were washed with PBS supplemented with 0.025 % Tween 20 and blocked with 1X ELISA/ELISpot Diluent (eBioscience). Goat ani-human IgG HRP (1:5000, Southern Biotech) was used as secondary antibody. ELISA signal was developed with Tetramethylbenzidine (eBioscience) and stopped with 1 M H_2_SO_4._ Optical Density (OD) was measured at 450 nm.

### HLA typing

Genomic DNA was isolated from PBMC of patients or healthy donor controls using the QIAmp DNA Blood Mini Kit (Qiagen) and submitted to DKMZ Germany for high-resolution HLA-typing. For patients HLA typing was confirmed at the expression level using arcasHLA to analyse single cell RNAseq data^46^.

## Data availability

Longitudinal TCR beta repertoire sequencing data will be available at 10.21417/TB2023NM. scVDJ and scRNA will be available at the Short Read Archive under BioProject PRJNA958150.

## Acknowledgements

We would like to express our gratitude to the patient and their relatives as well as to the clinical units. This study was supported by BWST_ISF2018-046 from the Baden-Württemberg-Stiftung to MP and the Dr Rolf M. Schwiete Foundation (2021-009) to LB and MP, as well as BioMed X funding to JML from the Janssen Pharmaceutical Companies of Johnson and Johnson. MP and KS are supported by the ‘INTERCEPT-H3’ grant from German cancer aid as well as the ‘Precision immunotherapy of brain tumors’ grant from the German Ministry of Education and NCT. TB, EG, IP and MP are supported by exploratory grants from Helmholtz-Institute for Translational Oncology Mainz (HI-TRON Mainz). EG is supported by technology development grants from the Federal Ministry of Education and Research (BMBF) and the Ministry of Science Baden-Württemberg within the framework of the Excellence Strategy of the Federal and State Governments of Germany. KL is funded by the Helmholtz International Graduate School (HIGS). KS is supported by a grant from the Hertie Network of Excellence in Clinical Neuroscience. We acknowledge the support of the DKFZ Genomics and Proteomics as well as of the Cytometry Core Facility, and that the schematics in Figure 1 and Extended Data Figure 3 were created using BioRender.com. Finally, we thank the DKFZ immune monitoring team for technical support, and Dr. Nathan Felix and Prof. Nina Papavasiliou for their valuable input throughout the project duration.

## Extended Data

**Extended Data Figure 1:**
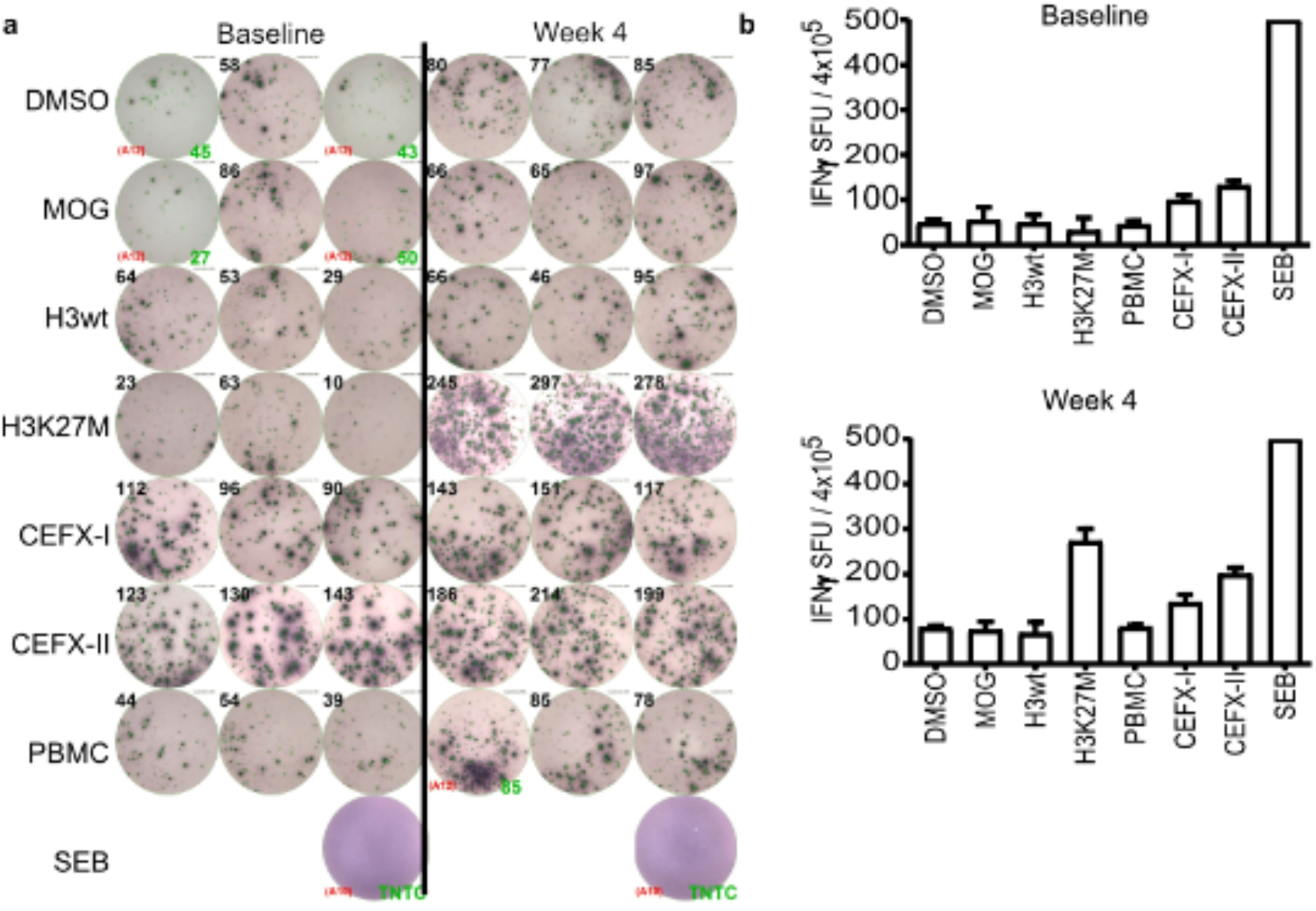
ELISpot at Baseline and Week 4 after treatment initiation of ID01. **a**, Spot counts of IFNγ ELISpot assay and their quantification (**b**) is shown. DMSO, Myelin oligodendrocyte glycoprotein (MOG) and only medium serve as negative controls. HLA class I (CEFX-1) and HLA class II (CEFX-II) stimulating peptide pools as well as Staphylococcal enterotoxin B (SEB) are used as positive controls. TNTC (=too numerous to count).

**Extended Data Figure 2:**
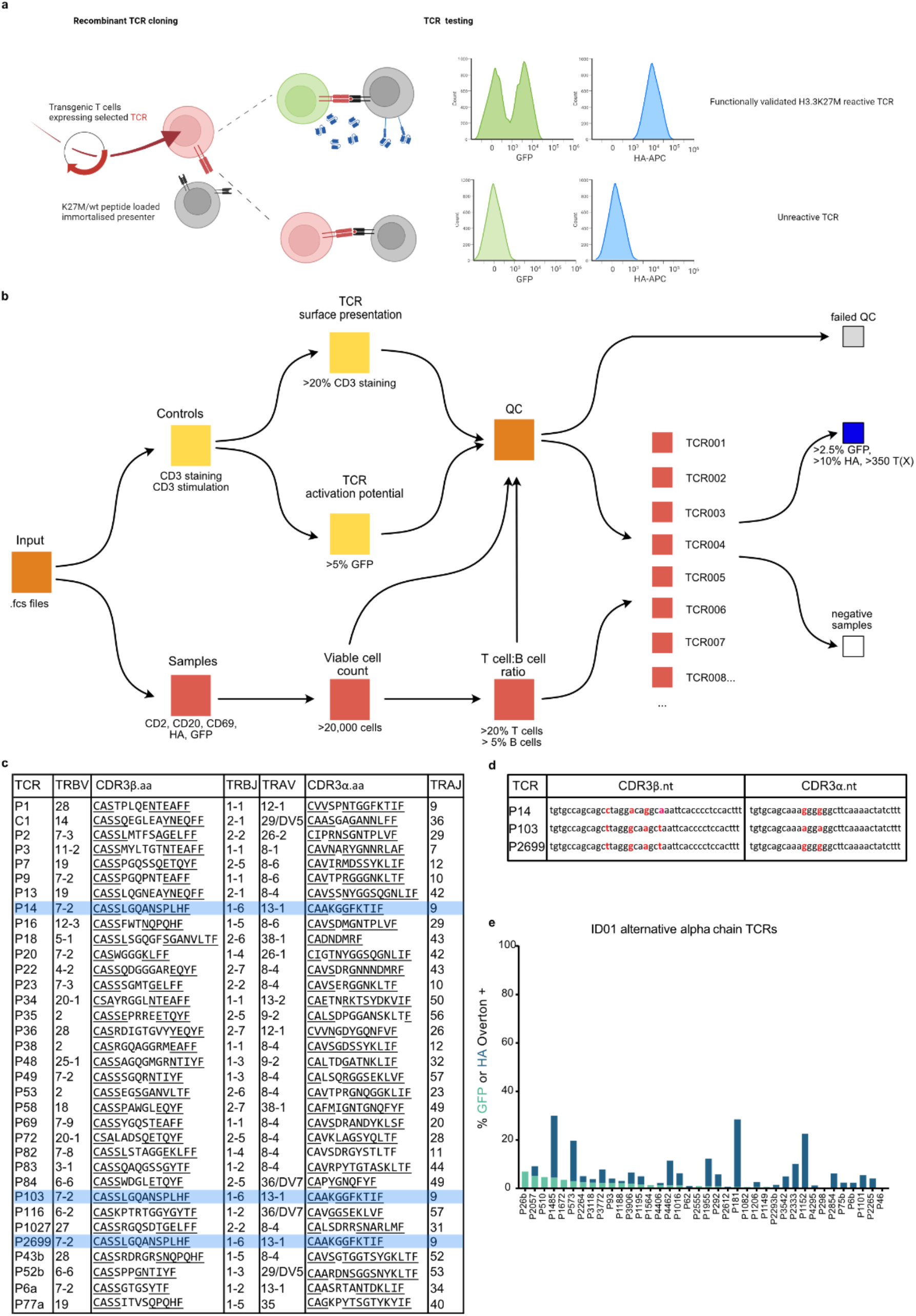
Screening platform and identification of H3K27M reactive TCRs. **a**, Schematic overview of T-FINDER platform. Selected TCRs are cloned and introduced into the Jurkat reporter T cell line and subsequently screened for reactivity in co-cultures with peptide-loaded autologous B-LCLs. Reactive TCR will lead to a dual reporter activation with a positive GFP signal in Jurkats cells and a scFv αCD19 mediated anti-HA signal on B-LCLs (upper panel). **b**, Scheme of workflow for identification of positive responding TCRs. Each TCR construct is first assessed for CD3 staining and CD3 stimulation. Only co-cultures that pass the indicated QC requirements will be considered for subsequent analyses. A TCR is considered reactive when both reporter signals reached the indicated thresholds (blue square). **c**, List of V and J gene segment usage and CDR3 amino acid sequences of positive TCRs. Underlined sequences are encoded by the V and J segments, respectively. **d**, Table highlighting nucleotide differences in the recombination junction sites of the convergently selected TCRs P14, P103, and P2699. **e**, Bar plot showing GFP and HA signals of alternative lower abundance alpha chains paired with beta chains shown in Fig 2a.

**Extended Data Figure 3:**
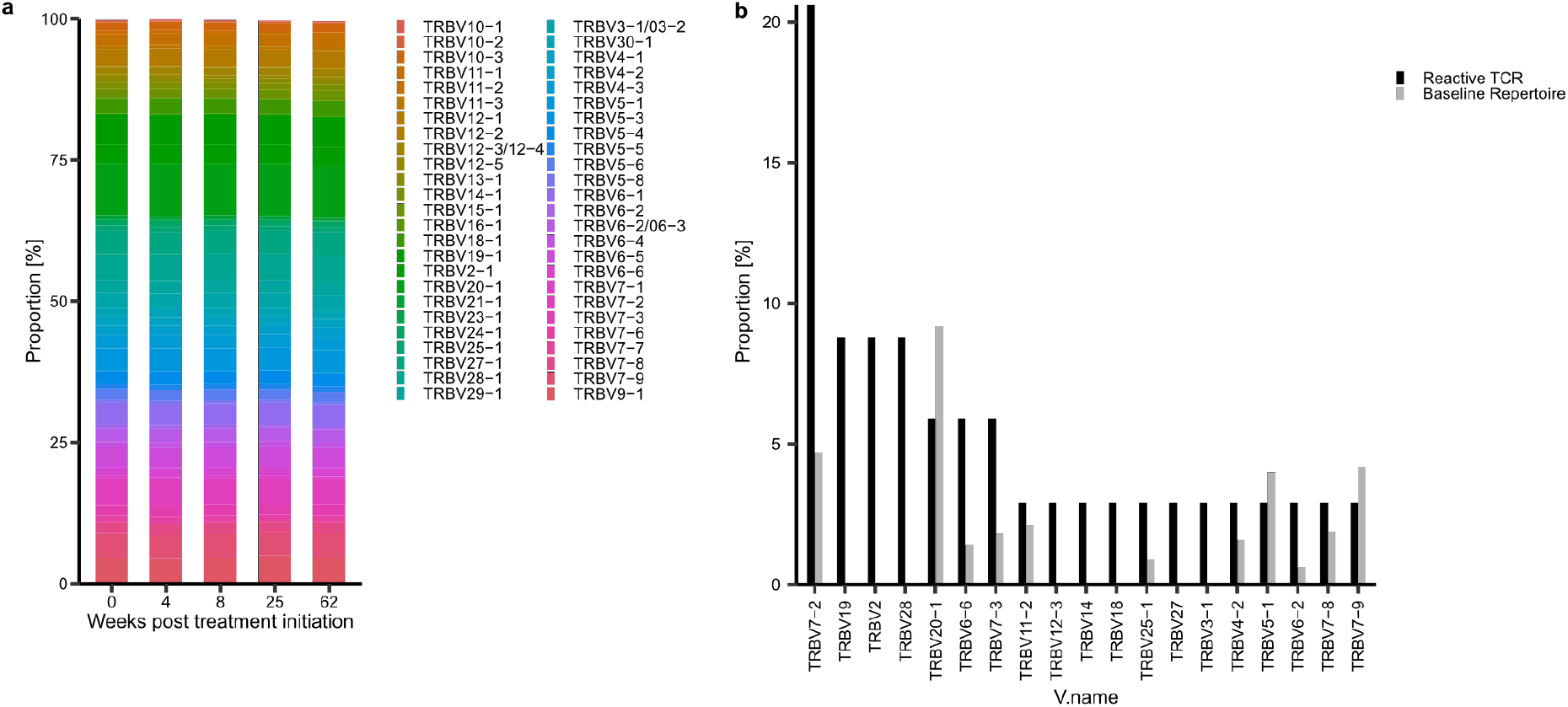
TRBV gene usage in patient ID01. **a**, Stacked bar chart showing TRBV gene usage and their proportion at indicated weeks. **b**, Bar charts depicting proportion of used TRBV genes in reactive TCRs and in baseline TCR repertoire.

**Extended Data Figure 4:**
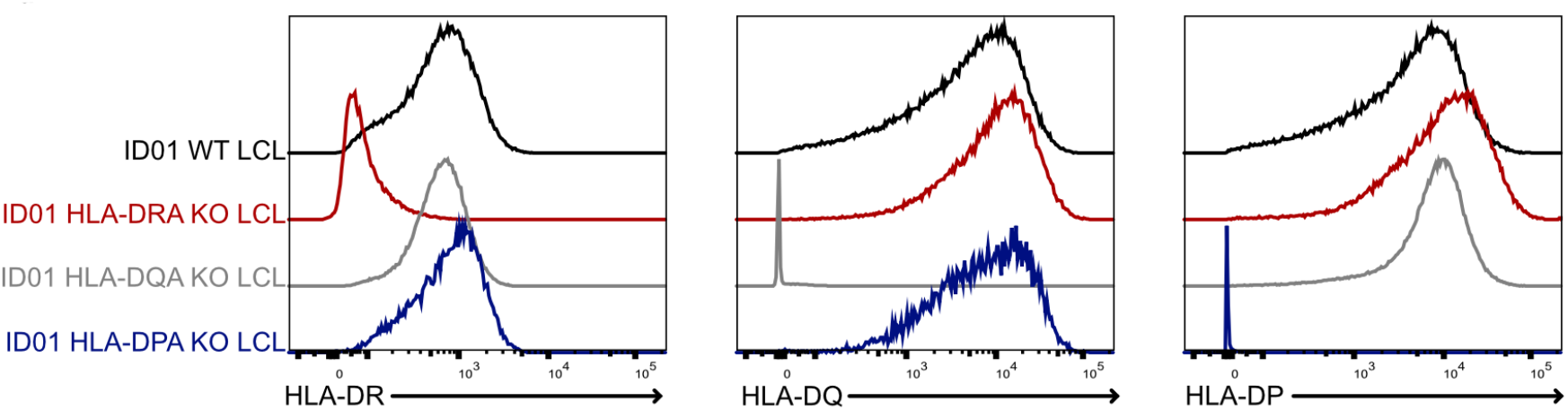
Validation of CRISPR/Cas9 mediated HLA class II knockouts. Histograms of generated HLA class II knockout lines of ID01 were verified by staining for HLA-DR, -DQ and –DP.

**Extended Data Figure 5:**
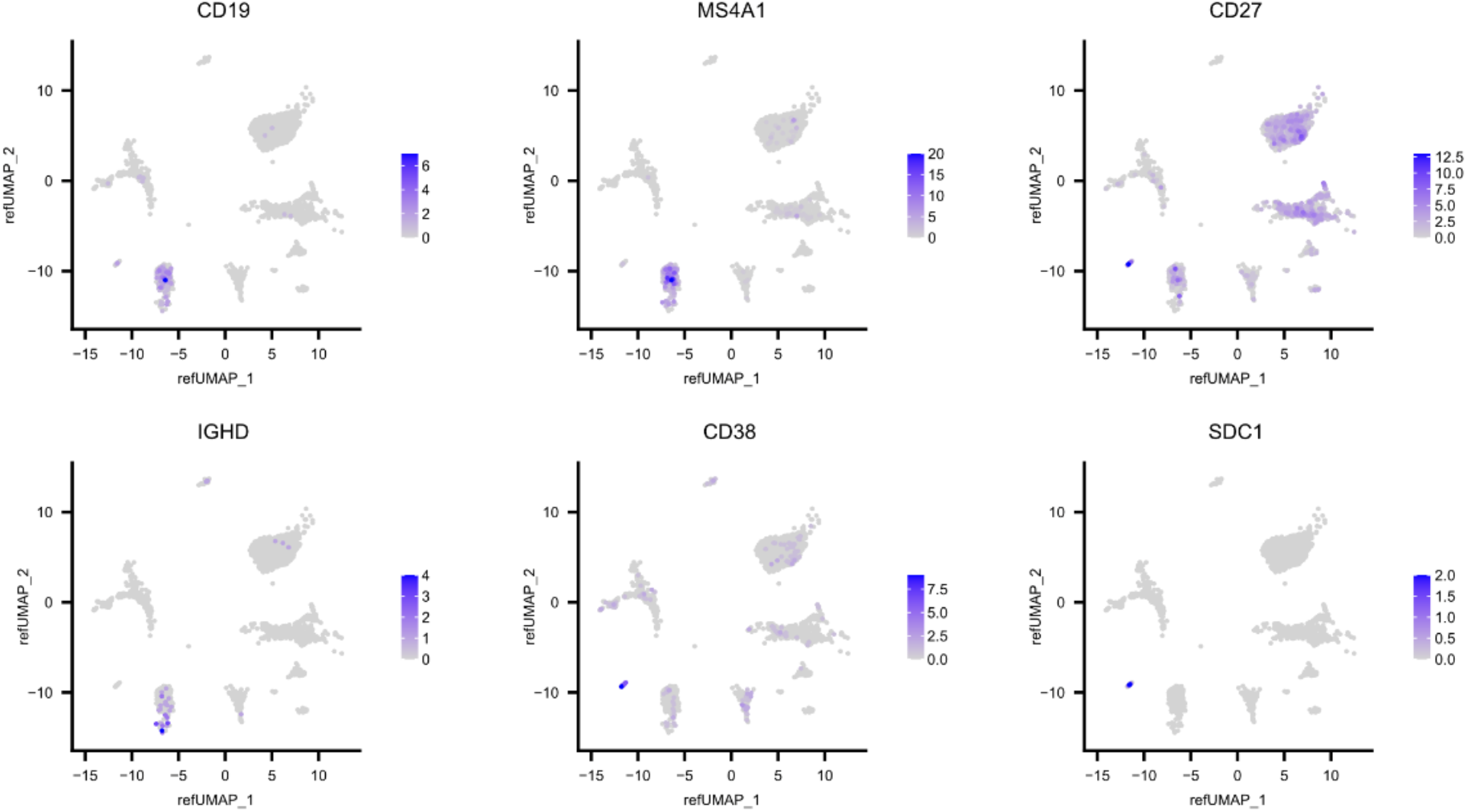
Gene expression of B cell specific markers in CSF of ID01. Feature Plots showing expression levels of relevant markers to identify subpopulations of B cells in Patient ID01.

